# An N-acetyltransferase required for EsxA N-terminal protein acetylation and virulence in *Mycobacterium marinum*

**DOI:** 10.1101/2023.03.14.532585

**Authors:** Owen A. Collars, Bradley S. Jones, Daniel D. Hu, Simon D. Weaver, Matthew M. Champion, Patricia A. Champion

## Abstract

N-terminal protein acetylation is a ubiquitous post-translational modification that broadly impacts diverse cellular processes in higher organisms. Bacterial proteins are also N-terminally acetylated, but the mechanisms and consequences of this modification in bacteria are poorly understood. We previously quantified widespread N-terminal protein acetylation in pathogenic mycobacteria (C. R. Thompson, M. M. Champion, and P.A. Champion, J Proteome Res 17(9): 3246-3258, 2018, https://doi:10.1021/acs.jproteome.8b00373). The major virulence factor EsxA (ESAT-6, Early secreted antigen, 6kDa) was one of the first N-terminally acetylated proteins identified in bacteria. EsxA is conserved in mycobacterial pathogens, including *Mycobacterium tuberculosis* and *Mycobacterium marinum*, a non-tubercular mycobacterial species that causes tuberculosis-like disease in ectotherms. However, enzyme responsible for EsxA N-terminal acetylation has been elusive. Here, we used genetics, molecular biology, and mass-spectroscopy based proteomics to demonstrate that MMAR_1839 (renamed Emp1, ESX-1 modifying protein, 1) is the putative N-acetyl transferase (NAT) solely responsible for EsxA acetylation in *Mycobacterium marinum*. We demonstrated that *ERD_3144*, the orthologous gene in *M. tuberculosis* Erdman, is functionally equivalent to Emp1. We identified at least 22 additional proteins that require Emp1 for acetylation, demonstrating that this putative NAT is not dedicated to EsxA. Finally, we showed that loss of *emp1* resulted in a significant reduction in the ability of *M. marinum* to cause macrophage cytolysis. Collectively, this study identified a NAT required for N-terminal acetylation in *Mycobacterium* and provided insight into the requirement of N-terminal acetylation of EsxA and other proteins in mycobacterial virulence in the macrophage.

**Significance Statement:** N-terminal acetylation is a protein modification that broadly impacts basic cellular function, protein turnover and disease in higher organisms. In bacteria, very little is understood how N-terminal acetylation impacts bacterial physiology and pathogenesis. Mycobacterial pathogens cause acute and chronic diseases in humans and in animals. ∼15% of mycobacterial proteins are N-terminally acetylated, but the enzymes responsible for this protein modification are largely unknown. We identified a conserved mycobacterial protein, MMAR_1839, that is required for the N-terminal acetylation of 23 mycobacterial proteins including EsxA, a protein essential for mycobacteria to cause disease. Loss of this enzyme from *Mycobacterium marinum* reduced macrophage killing, which is required for bacterial spread in the host. Defining the acetyltransferases responsible for the N-terminal protein acetylation of essential virulence factors could lead to new targets for therapeutics against mycobacterial pathogens.

## Introduction

The ESX-1 (ESAT-6 system-1) protein secretion system is essential for mycobacterial pathogenesis. Early during macrophage infection, ESX-1 is required for damaging the phagosomal membrane (1-5), allowing mycobacterial pathogens to access the macrophage cytosol (6, 7). The exposure of the *Mycobacterium* and its secreted factors to the cytoplasm combats the host response, and causes macrophage cytolysis (4, 8-10). *Mycobacterium* lacking the ESX-1 system are retained in the phagosome and attenuated (6). EsxA (ESAT-6) is a major mycobacterial virulence factor that is required for the pathogenesis of *M. tuberculosis* and other mycobacterial pathogens (3, 11-13). EsxA is secreted by ESX-1 and is required for the secretion of the majority of the ESX-1 substrates (1, 14, 15). EsxA may play additional roles in the host downstream of phagosomal lysis (16-22).

EsxA was one of the first bacterial proteins recognized to be N-terminally acetylated (23). EsxA forms a heterodimer with EsxB, another secreted ESX-1 component (1, 24). The acetylation state of the EsxA N-terminus was reported to mediate interaction with EsxB *in vitro* (23, 25). We previously reported an inverse correlation between EsxA acetylation and virulence (26). Aguilera et al mutated the 2^nd^ residue of EsxA to abrogate N-terminal acetylation. Their study suggested that N-terminal acetylation of EsxA was required for Esx1-mediated phagosomal lysis and macrophage cytolysis by *M. marinum* (25). *M. marinum* is an established model for studying the mycobacterial ESX-1 system (27). Importantly, deletion of *esx-1* genes in *M. marinum* is functionally complemented by the expression of orthologous genes from *M. tuberculosis*, demonstrating that the two systems share a conserved function (28).

N-terminal acetylation is the covalent addition of an acetyl group to the α amino group of the N-terminal amino acid of a protein by N-acetyltransferases (NATs) (29-31). NATs can irreversibly acetylate the iMet (initiator Met, following deformylation) or the first amino acid following iMet cleavage (32, 33). In higher organisms, including humans, yeasts and plants, ∼65-85% of proteins are N-terminally acetylated by 7 NATs (29). In higher organisms, N-terminal acetylation directly impacts protein function through a variety of mechanisms (29, 34-42). In contrast, ∼10-15% of bacterial proteins are N-terminally acetylated (30, 43-46). Using quantitative N-terminomics, we observed that ∼11 and ∼15% of proteins in *M. tuberculosis* and *M. marinum*, respectively, are N-terminally acetylated during standard laboratory growth *in vitro* (43). Our previous work revealed that in addition to EsxA, several additional ESX-1 substrates and at least one ESX-1 membrane component are also N-terminally acetylated (43).

Bacterial genomes encode several putative NATs, which are part of the GNAT (GCN-5 related N-acetyltransferase) family (30, 47). It is not possible to predict if the putative NAT acetylates Lys residues (KATs), small molecules, antibiotics, and/or protein N-termini (30). There are 27 predicted NATs in *M. tuberculosis*, 23 of which are conserved in *M. marinum* (43). The best characterized NAT is RimI, which N-terminally acetylates the S18 rRNA protein and functions broadly as a generalist KAT in *Escherichia coli, Salmonella* and in *Mycobacterium* (47-51). The individual NATs responsible for N-terminal acetylation of specific mycobacterial proteins are lacking, limiting our understanding of the role of this modification in mycobacterial virulence and physiology.

The EsxA NAT has remained elusive, and it is unknown if one or more NATs contribute to EsxA acetylation. Changes in EsxA acetylation are difficult to assess because the amino acid composition of EsxA yields a large N-terminal tryptic fragment with poor chromatographic performance by mass-spectroscopy based proteomics (52-54). Top-down approaches do not rapidly assign all N-terminal isoforms of EsxA (52-54). In this study, we sought to identify the NAT responsible for N-terminally acetylating

EsxA to further understand N-terminal acetylation in *Mycobacterium*. Based on the conservation of EsxA and the putative NATs between *M. marinum* and *M. tuberculosis*, we hypothesized that we could leverage the use of *M. marinum* to identify the conserved EsxA NAT. To test this hypothesis, we used an N-terminal acetyl-EsxA antibody coupled with a knockout *M. marinum* strain collection to identify the EsxA NAT. We measured EsxA acetylation in the presence and absence of the putative NAT using western blot analysis, MALDI, and label-free quantitative mass spectrometry (LFQ). We tested ESX-1 function and mycobacterial virulence using *in vitro* systems and macrophage model of infection.

## Results

### In vitro *discrimination between the EsxA and acetyl-EsxA N-termini*

We hypothesized that an antibody specific to the acetylated N-terminus of EsxA could be used a tool to identify the NAT(s) responsible for Nt-acetylation of EsxA. We obtained and characterized a polyclonal antibody synthesized against an acetylated N-terminal EsxA peptide (Fig. 1A, Ac-EsxA). We performed a dot blot to determine if the Ac-EsxA antibody specifically recognized the acetylated N-terminus of EsxA. As shown in Figure 1B, the Ac-EsxA antibody specifically produced signal where the Ac-EsxA peptide was spotted on the nitrocellulose (red) but not where the unacetylated peptide was spotted (outline). In contrast, the commercially available EsxA antibody raised against the same unacetylated peptide specifically produced signal where the unacetylated EsxA peptide was spotted onto the nitrocellulose (green), but not where the acetylated peptide was spotted. From these data, we conclude that the Ac-EsxA antibody discriminates between the acetylated and unacetylated forms of the EsxA N-terminal peptide. Moreover, the commercial EsxA antibody specifically recognizes the unacetylated form of EsxA.

**Figure 1.**
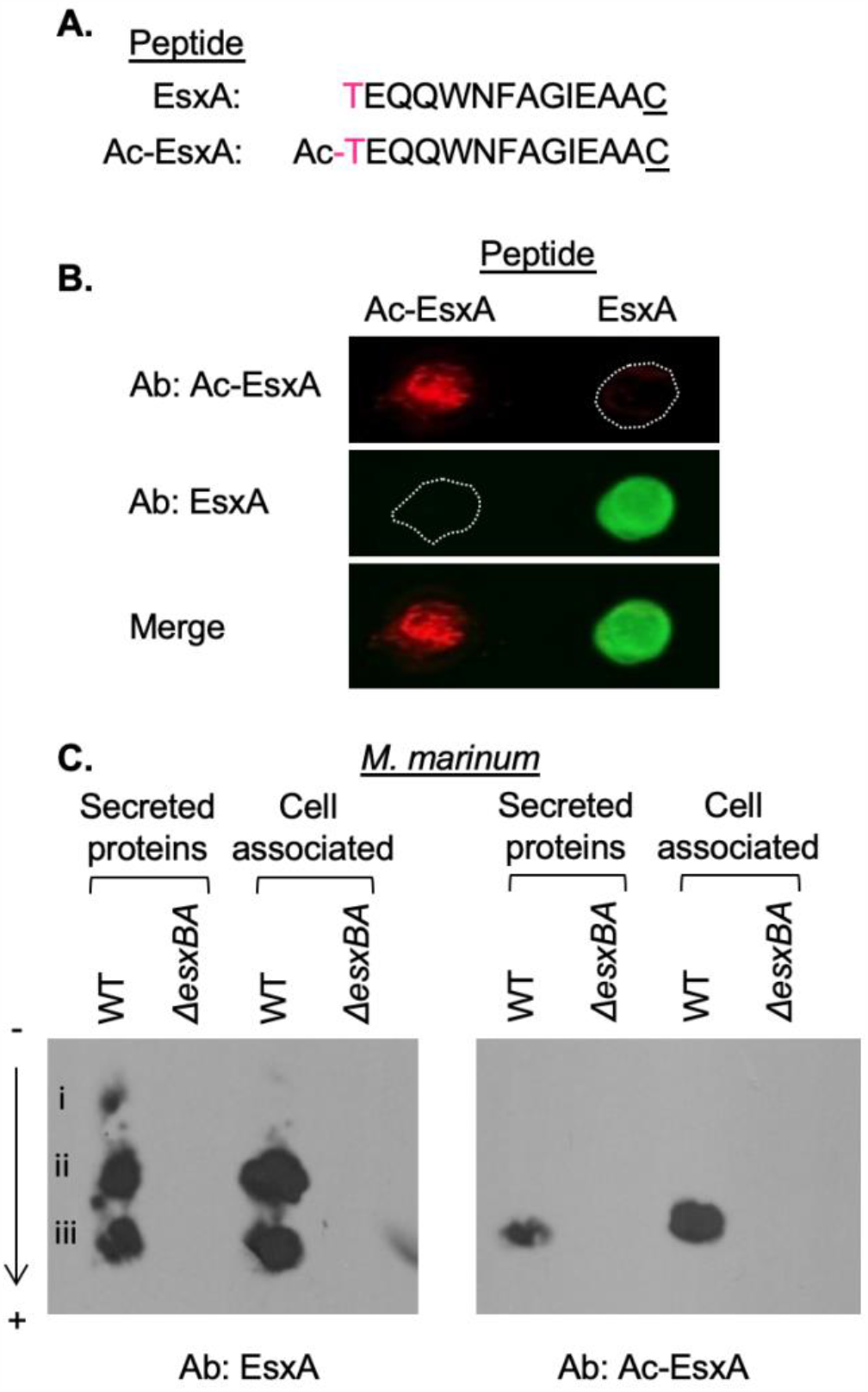
The Ac-EsxA antibody specifically recognizes EsxA in complex mixtures. **A**. The EsxA and Ac-EsxA N-terminal peptides. The “C” is not native to the EsxA protein **B**. Dot blot of EsxA N-terminal peptides. 20μg of each peptide were spotted on nitrocellulose and immunoblotted with the αEsxA and αAc-EsxA antibodies. The image is representative of three independent replicates **C**. NUT-PAGE of secreted and cell associated proteins from WT and Δ*esxBA M. marinum* strains. 20μg of protein was loaded in each lane. The image is representative of at least three independent biological replicates.

We next tested if the Ac-EsxA antibody could detect the acetylated version of the EsxA protein in a complex mixture of proteins. We collected cell-associated and secreted protein fractions from the wild-type (WT) and Δ*esxBA M. marinum* strains. We separated the proteins by charge using the neutral pH urea Triton polyacrylamide gel electrophoresis (NUT-PAGE) system, which can separate acetylated and unacetylated proteins (55). As shown in Figure 1C, NUT-PAGE followed by western blot analysis allowed for the separation of several protein species (i-iii) detected by the EsxA antibody in the protein fractions collected from the WT *M. marinum* strain. All three of these species were absent from the protein fractions generated from the Δ*esxBA* strain, which fails to produce EsxA protein. Notably, only species iii, which is the most negatively charged EsxA species, was reliably detected by the Ac-EsxA antibody. From these data we conclude that we can separate and detect acetylated EsxA from a complex mixture of *M. marinum* proteins.

### *N-terminal acetylation of EsxA and other proteins is dependent on* MMAR_1839

The EsxA proteins from *M. marinum* and *M. tuberculosis* are identical through the 15^th^ amino acid, and 92% identical overall (Figure S1). Therefore, we reasoned that the NAT responsible for acetylating EsxA would be highly conserved between *M. marinum* and *M. tuberculosis*. We identified the five proteins with predicted GNAT domains that were the most highly conserved between the two species (Table 1). Using allelic exchange, we generated unmarked deletions of each putative NAT gene in *M. marinum*. We confirmed the deletion of each gene by PCR (Fig. S2) and targeted DNA sequencing.

**Table 1:** The top five conserved putative NATs between *M. marinum* and *M. tuberculosis*. Sequences were obtained from Mycobrowser. The % protein identity was determined using Protein BLAST.

| *M. marinum*<br>gene | *M. tuberculosis*<br>gene | % Identity<br>(Protein) |
| --- | --- | --- |
| *MMAR_1067* | *Rv0730* | 90.9% |
| *MMAR_1968* | *argA* | 87.3% |
| *MMAR_4519* | *rimJ* | 87.56% |
| *MMAR_1839* | *Rv2867* | 87.32% |
| *MMAR_1882* | *elaA* | 86.75% |

We hypothesized that we would not detect acetylated EsxA from *M. marinum* strains lacking an EsxA specific NAT. We collected cell-associated proteins from *M. marinum* strains lacking each of the five most conserved NATs. We measured EsxA and acetylated-EsxA in these strains using western blot analysis, as compared to proteins generated from the WT and *ΔesxBA* strains. As shown in Figure 2A, both the unacetylated and acetylated EsxA proteins were detected in lysates generated from the WT *M. marinum* strain (lane 1). These data are consistent with prior studies demonstrating that both species exist in the WT strain (23, 26, 43, 56-58). Both EsxA species were lacking from the Δ*esxBA* strain (lane 2), demonstrating the specificity of both antibodies to EsxA. Acetylated and unacetylated EsxA were present in the lysates generated from the Δ*MMAR_1067*, Δ*MMAR_4519* or Δ*MMAR_1882* strains (lanes 3, 5 and 7). The Δ*MMAR_1968* strain lacks the ArgA NAT and is auxotrophic for arginine (59). Addition of L-arginine to the growth media allowed detection of acetylated and unacetylated EsxA from the Δ*MMAR_1968* lysates (Fig. 2B). Deletion of the *MMAR_1839* gene resulted in detection of the EsxA protein (Fig. 2A, lane 6), but not the Ac-EsxA protein from this lysate. From these data we conclude that MMAR_1839 is required for EsxA acetylation in *M. marinum*. We renamed *MMAR_1839*, ESX-1 modifying protein-1 (Emp1).

**Figure 2.**
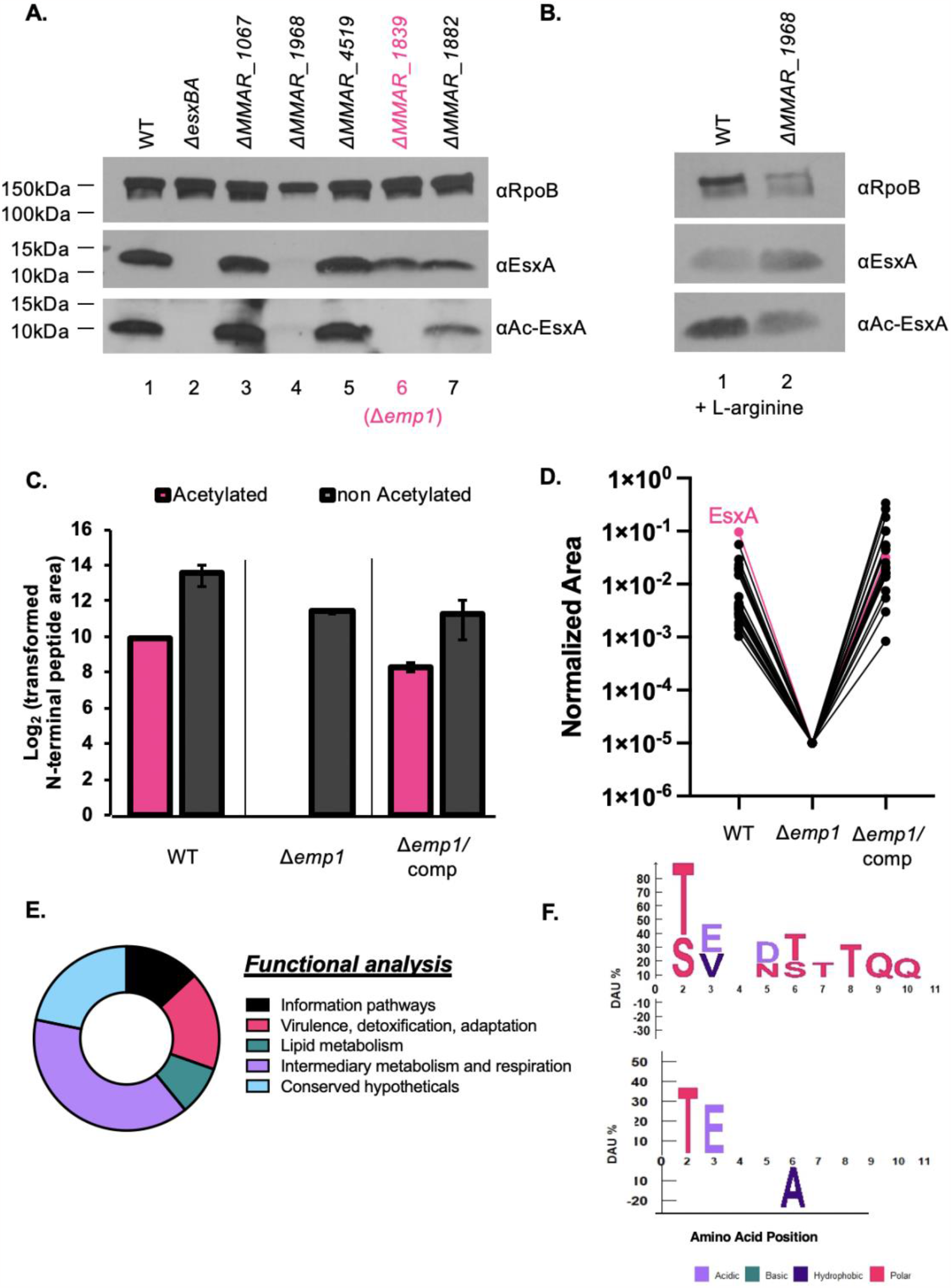
MMAR_1839 (Emp1) is required for the acetylation of EsxA and other proteins. **A**. and **B**. Western blot analysis of cell-associated proteins from the indicated *M. marinum* strains. 10μg of protein was loaded in each lane. In B, 2μM of L-arginine was added to the culture media. Both images are representative of at least three biological replicates. RpoB is a control for loading. **C**. MS Analysis of relative abundance of acetylated and non-acetylated N-terminus of EsxA. Label Free Quantitative (LFQ) proteomics intensity of the EsxA N-terminal peptide from WT, Δ*emp1*, Δ*emp1/*complemented strains. Normalized intensity was transformed by 10^4^ to convert the Log2 values to positive integers. Propagated error was performed on technical triplicates. **D**. K-means clustering of all N-terminally acetylated proteins observed from bottom-up proteomics in WT, Δ*emp1*, Δ*emp1/*complemented strains. Shown is the cluster that contained EsxA. **E**. Functional analysis from Mycobrowser of the 22 proteins that clustered with EsxA from the k-means analysis from *M. marinum*. For conserved hypothetical proteins, if the closest ortholog in *M. tuberculosis* was annotated, that annotation was used instead. **F**. ICE Logo from the protein N-termini in D and E. Differential Amino Acid Usage (DAU) tests were used to determine overrepresented and underrepresented amino acids at specific N-terminal amino acid positions (77). Fisher’s exact test with a significance scoring of *P*<0.05 was used to determine significance. The top logo is the sequence of these protein N-termini compared to the whole *M. marinum* proteome. The bottom logo is the same proteins N-termini compared to the *M. marinum* N-terminome from (See Supplemental Material) (43). All R code is available on GitHub (https://github.com/Champion-Lab/ESXA_Acetylation) along with a list of data analysis steps.

To determine if Emp1 was required for the acetylation of additional mycobacterial proteins, we performed label-free quantitative (LFQ) mass spectrometry to measure the relative changes in acetylation and protein levels in the Δ*emp1* strain as compared to the WT and complemented strains (Dataset S1, Raw and Trimmed Data tabs S1A and S2B). To confirm the western blot analysis, we first compared the levels of EsxA and Ac-EsxA from the three strains. As shown in Figure 2C, the levels of EsxA were comparable in all three strains (grey bars, Dataset S1, tab S1C). While we detected Ac-EsxA in the WT and the complemented strains, we did not detect Ac-EsxA in the Δ*emp1* strain. From these data, we conclude that Emp1 is the only EsxA NAT in *M. marinum*.

We next performed *k*-means (60) clustering analysis for each N-terminally acetylated protein in the dataset, using the biological replicate with the best coverage (Dataset S1, tab S1C). Using this approach, we systematically identified patterns between the WT, Δ*emp1*, and complementation strains across every protein. The variables considered for the clustering were the LFQ area ratios from the following strains: Δ*emp1*/complement, Δ*emp1*/WT, and complement/WT. The proteins were clustered into three groups, using 25 random starting points. We reasoned that proteins that clustered with EsxA were potential acetylation targets of Emp1, as they also exhibited loss of acetylated intensity in Δ*emp1*, which was restored upon complementation. The acetylation intensity patterns of proteins identified from the clustering were compared across all biological replicates.

Using this approach, we identified a cluster of proteins whose N-terminal acetylation followed a similar pattern to the levels of acetylated EsxA. 23 proteins, including EsxA, exhibited undetectable levels of N-terminal acetylation in the Δ*emp1* strain, and restoration of acetylation in the complemented strain, similar to the WT strain (Fig. 2E, EsxA highlighted in pink, Dataset tab S1E). Functional analysis revealed that the majority of proteins that depend on Emp1 for acetylation are predicted to function in lipid metabolism or intermediary metabolism and respiration (Fig. 2E). Four proteins, including EsxA, are involved in virulence. More than half of the potential protein targets are annotated as essential *in vitro* in *M tuberculosis*. Finally, we analyzed the N-terminal amino acid sequence of the proteins dependent upon Emp1 for N-terminal acetylation. When comparing the N-terminal sequences of the putative targets of Emp1 against the entire *M. marinum* proteome, we a see a strong negative bias for basic residues within these first ten amino acids (Fig. 2F, upper). This is likely due to the use of trypsin for the mass-spectrometry based proteomics, which cleaves after Lys and Arg. Consequently, those peptides are underrepresented in the first 10 amino acids of the N-termini as they would not be observed due to their small size (43, 54). There was also a strong preference for threonine, with mild preference for serine at the second amino acid position, consistent with our prior work (43). Comparison of the N-terminal sequences of the putative targets of Emp1 against the *M. marinum* N-terminal acetylome (43), a strong preference for threonine and glutamic acid at the second and third amino acid positions, and a significant underrepresentation of alanine at the sixth amino acid position (Fig. 2F, lower). Together, these data demonstrate that Emp1 is required for the N-terminal acetylation of EsxA and at least 22 other proteins in *M. marinum*.

### Emp1, and therefore EsxA acetylation, is dispensable for EsxA/EsxB secretion from M. marinum

The identification of Emp1 allowed us to test the role of EsxA acetylation on ESX-1 function in the presence of the wild-type *esxA* gene in *M. marinum*. It was previously suggested that EsxA acetylation impacted the interaction between EsxA and its binding partner, EsxB (23). The EsxA-EsxB interaction is required ESX-1 function; EsxA-EsxB interaction is required for protein stability and for targeting the EsxA-B pair for ESX-1 (1, 56, 61). If N-terminal acetylation of EsxA was required for interaction between EsxA and EsxB, then we would expect a loss of EsxB protein and a corresponding loss of EsxA and EsxB secretion from the Δ*emp1* strain. We generated cell-associated and secreted protein fractions from *M. marinum* strains. As shown in Figure 3A, EsxA, Ac-EsxA and EsxB were produced (lane 1) and secreted (lane 7) from the wild-type *M. marinum* strain. Deletion of the *eccCb*_*1*_ gene resulted in reduced levels of EsxA and Ac-EsxA (lane 2), consistent with the reduced levels of the EsxA substrate in the absence of secretion (1, 62). Neither EsxA, Ac-EsxA nor EsxB were secreted from the *ΔeccCb*_*1*_ strain (lane 8). Although EsxA and EsxB were produced in the Δ*emp1* strain, Ac-EsxA was not detected (lane 3). Both EsxA and EsxB were secreted from the Δ*emp1* strain (lane 9). Constitutive expression of the *emp1* gene restored the production (lane 4) and secretion of Ac-EsxA (lane 10). Likewise, expression of the orthologous gene from *M. tuberculosis* Erdman (*ERD_3044*) restored the production and secretion of Ac-EsxA from the Δ*emp1* strain (lanes 5 and 11). Finally, we mutated the predicted active site of Emp1 (W223A), which would render the enzyme unable to bind Ac-CoA (63). Expression of *emp1W223A* in the Δ*emp1* strain did not restore production or secretion of Ac-EsxA (lanes 6 and 12). Together, these data demonstrate that Emp1 is required for the acetylation of EsxA. Contrary to existing models (25), the loss of EsxA acetylation did not result in a loss of the EsxA or EsxB protein, suggesting that the acetylation state of EsxA is dispensable for EsxA/EsxB interaction and secretion from *M. marinum* during *in vitro* growth. Our data supports that Emp1 likely functions as an NAT in *M. marinum*, and is functionally conserved in *M. tuberculosis*.

**Figure 3.**
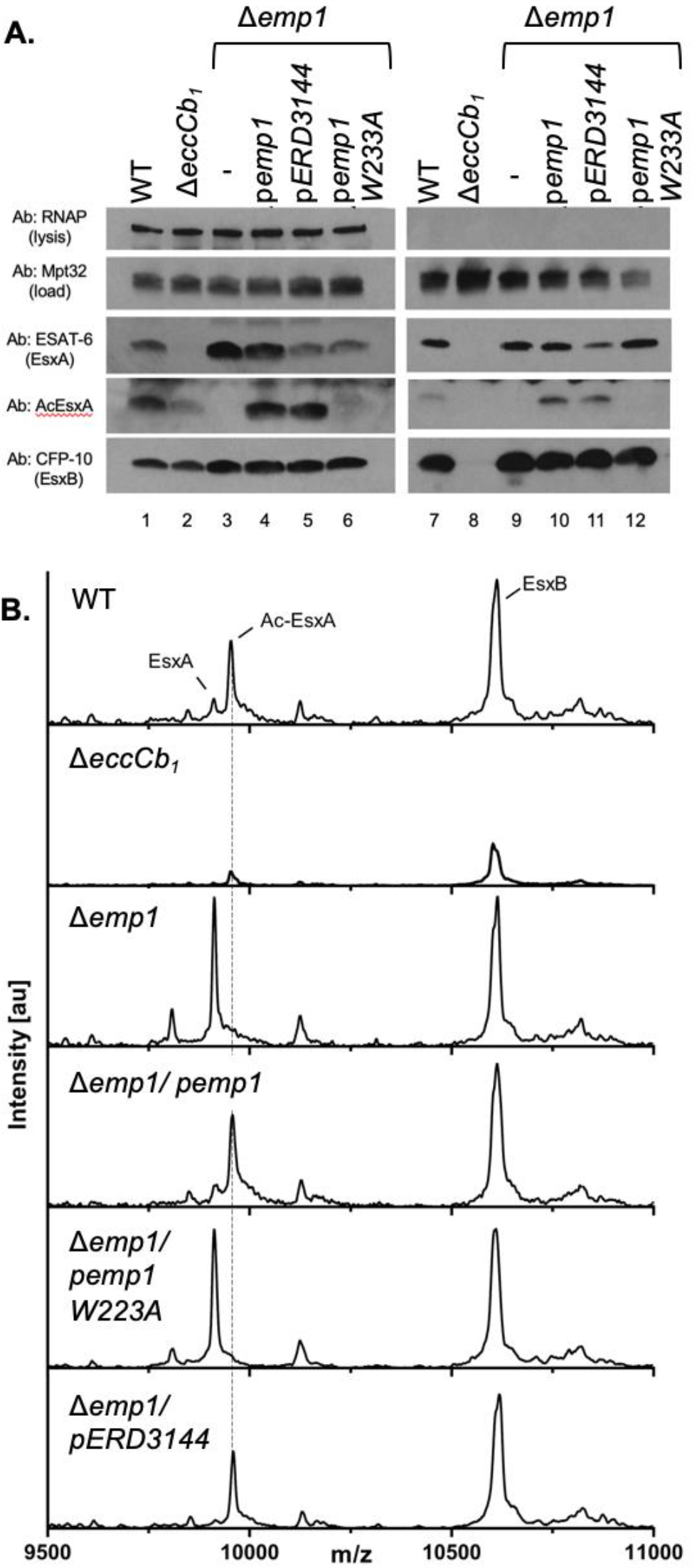
Emp1 is dispensable for EsxA and EsxB stability and secretion from *M. marinum*. **A**. Western blot analysis of cell-associated and secreted proteins from *M. marinum* in the presence and absence of Emp1 **B**. whole colony MALDI-TOF MS. Spectra generated by whole colony MALDI-TOF for wild-type and mutant and complemented *M. marinum* strains are shown. The labeled peaks correspond to EsxA (9915 m/z), acetylated EsxA (9957 m/z), and EsxB(10,606 m/z Da), respectively. The dotted line was added for clarity.

We sought a more sensitive approach to confirm that Emp1 was required for acetylation of EsxA in *M. marinum*. Because there are 22 additional putative conserved NATs in encoded in the *M. marinum* genome, we wanted to further verify that EsxA was completely unacetylated in the Δ*emp1* strain and rule out cross-talk by other putative NATs in the absence of *emp1*.

We previously demonstrated that both acetylated and unacetylated EsxA are resolved in proteins washed from the surface of *M. marinum* colonies using whole-colony MALDI-TOF-MS (58). Using this approach, we detected peaks consistent with both unacetylated (9915 m/z) and acetylated (9957 m/z) EsxA from surface associated proteins isolated from WT *M. marinum* colonies (Fig. 3B). We also detected surface associated EsxB (10,606 m/z). The Δ*eccCb*_*1*_ strain is a lysis control, because this strain produces but does not secrete EsxA and EsxB (1, 64). Both EsxA species and EsxB were significantly diminished from the proteins isolated from the surface of the Δ*eccCb*_*1*_ strain (1, 58). Therefore, the observed peaks are due to the secretion of EsxA and EsxB to the cell surface. Proteins isolated from the surface of the *Δemp1* strain resulted in a single EsxA peak which corresponded to the unacetylated EsxA protein, and a peak for EsxB. The acetylated EsxA peak was completely abrogated. Expression of the wild-type *emp1* gene, but not the *emp1W223A* gene, restored the peak corresponding to the Ac-EsxA protein. From these data we conclude that deletion of the *emp1* gene results in a complete loss of Ac-EsxA in *M. marinum*, demonstrating that Emp1 is solely responsible for the acetylation of EsxA *in vivo*. The absence of acetylation in the W223A active-site mutant of Emp-1 (Fig. 3B) demonstrates that functional Emp-1 is required for the acetylation of EsxA.

Moreover, our findings demonstrate that EsxA and EsxB are secreted from *M. marinum* independently of EsxA-N-terminal acetylation during *in vitro* growth.

### emp1 is dispensable for ESX-1 function but required for macrophage cytolysis

We next tested if *emp1* was required for *M. marinum* pathogenesis. It was previously reported that the N-terminal acetylation of EsxA was required for ESX-1-dependent phagosomal lysis (25). Hemolytic activity is one measurement of ESX-1 function *in vitro* (64, 65). *M. marinum* lyses red blood cells in a contact-dependent, ESX-1-dependent manner (64). EsxA is required for the hemolytic activity of *M. marinum*, likely because it is required for the secretion of the majority of the ESX-1 substrates (14, 15, 28, 66). Importantly, the *ΔesxA M. marinum* strain is non-hemolytic (14). Because the acetylation of EsxA depends on Emp1, we reasoned that if EsxA acetylation was required for EsxA function, the Δ*emp1* strain would have altered hemolytic activity.

As shown in Figure 4A, WT *M. marinum* lysed sheep RBCs (sRBCs), while the Δ*eccCb*_*1*_ strain (which fails to secrete ESX-1 substrates) exhibited significantly reduced hemolytic activity (*P*<.0001, relative to the WT strain). Water and PBS (cell-free) were used as positive and negative controls, respectively. The activity of the Δ*eccCb*_*1*_ strain was not significantly different from the PBS control (*P*<.9999). The hemolytic activities of the Δ*emp1* and the Δ*emp1* complemented strains were not significantly different from the WT strain (*P*=.4837 and *P*=.9998) or each other (*P*=.2689). From these data we conclude that Emp1 is dispensable for hemolytic activity of *M. marinum*. Because ESX-1 mediates hemolysis, the data suggest that the acetylation of EsxA is also dispensable for hemolysis, and are consistent with the secretion of EsxA and EsxB from the Δ*emp1* strain (Fig. 3B). Finally, because additional ESX-1 substrates required for hemolysis depend upon EsxA for secretion, our data suggest that the secretion of additional ESX-1 substrates occurs independently of EsxA acetylation.

**Figure 4.**
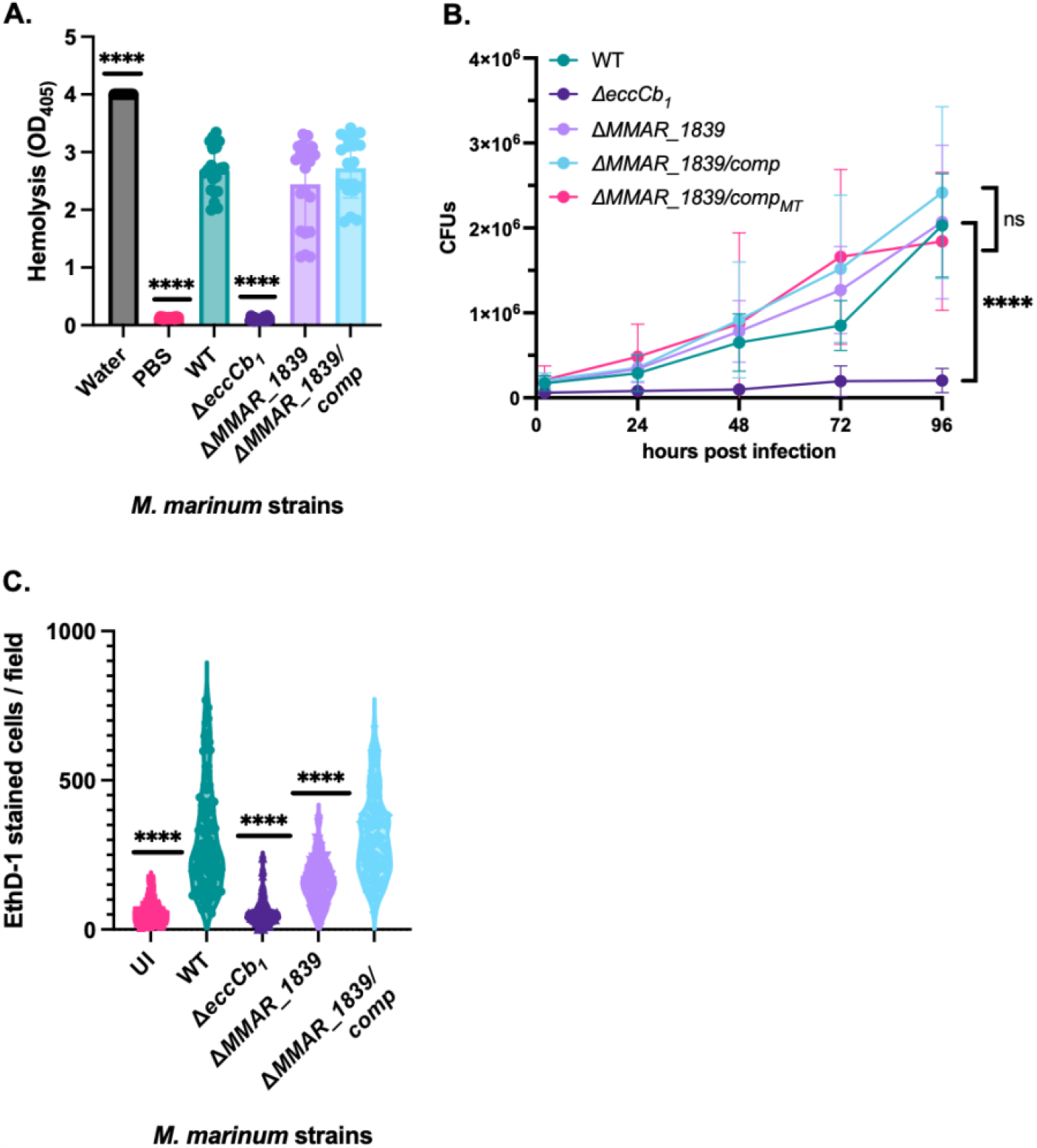
Emp1 is dispensable for hemolytic activity and growth in a macrophage model, but is required for cytolytic activity. **A**. Hemolytic activity of *M. marinum* strains. Data shown includes 7 biological replicates each in technical triplicate. Each data point is a technical replicate. Statistical analysis was performed using an ordinary one-way ANOVA (*P*<.0001) followed by a Tukey’s multiple comparison test. Significance shown is compared to the WT strain. Other important comparisons are discussed in the text. **B**. CFU analysis of *M. marinum* strains. MOI = 0.2 plated in triplicate, represents 3 biological replicates. Significance was determined using a 2-way RM ANOVA (*P*<.0001) followed by a Tukey’s multiple comparison test. Significance shown is compared to the WT strain, for the 96hpi. However, the CFUs from the Δ*eccCb*_*1*_ strain were significantly lower than the WT strain throughout the experiment. P values comparing Δ*eccCb*_*1*_ vs WT were as follows: 0 hpi ** *P*=.0035, 24 hpi, * *P*=.0162, 48 hpi *** *P* = .0001, 72 hpi and 96 hpi **** *P* <.0001. The WT strain was not significantly different from the additional strains at any time point. **C**. Cytolytic activity of *M. marinum* strains in RAW 264.7 cells, 24 hpi, MOI = 5. The data includes at least 3 biological replicates with 10 fields selected for each infection. Each data point is the number of red cells per field. Significance was determined using an ordinary one-way ANOVA (*P*<.0001) followed by a Sidak’s multiple comparison test. Significance shown is compared to the WT strain, with additional comparisons discussed in the text. **** *P*<.0001.

ESX-1 activity can also be measured during macrophage infection. We reasoned that if acetylation of EsxA was required for function, then the *Δemp1 M. marinum* strain would be attenuated for growth in a macrophage model of infection, similar to the Δ*eccCb*_*1*_ strain (1, 67, 68). We infected RAW 264.7 cells with *M. marinum* and measured colony forming units over time. As shown in Figure 4B, the WT *M. marinum* strain grew over time in the macrophages, while the Δ*eccCb*_*1*_ strain was attenuated for growth. Deletion of the *emp1* gene did not impact the ability of *M. marinum* to grow in the macrophage (*P*<.0001). Growth of the Δ*emp1* and the Δ*emp1* complemented strains was not significantly different from the WT strain. From these data, we conclude that *emp1* is dispensable for *M. marinum* growth in the macrophage. Moreover, it is unlikely that the function of the ESX-1 system, including EsxA, is impacted by N-terminal acetylation in this model of infection. Following ESX-1 dependent phagosomal lysis, *M. marinum* are released into the cytoplasm, promoting macrophage cytolysis through ESX-1-independent mechanisms (3, 6, 7, 69-72). We next tested if Emp1 was required for macrophage cytolysis. We infected RAW 264.7 cells with *M. marinum* and measured uptake of the membrane impermeable dye, Ethidium homodimer 1 (EthD-1). As shown in Figure 4C, infection of RAW 267.4 cells with wild-type *M. marinum* resulted in a significant level of cytolysis, as reflected by EthD-1 uptake, compared to the uninfected cells (*P*<.0001). The Δ*eccCb*_*1*_ strain exhibited significantly less cytolysis than the WT strain (*P*<.0001), similar to the uninfected control (*P*>.9999). Deletion of the *emp1* gene resulted in a significant reduction in cytolysis compared to the WT strain (*P*<.0001). The levels of EthD-1 uptake following infection with the Δ*emp1* strain was significantly higher than those following infection with the Δ*eccCb*_*1*_ strain and the uninfected control (*P*<.0001). Constitutive expression of the *emp1* gene in the Δ*emp1* strain restored cytolysis to levels similar to the WT strain. To confirm that these strains were not attenuated due to the spontaneous loss of the outer lipid PDIM, we performed TLC analysis. All of the *emp1* strains produced PDIM similar to the WT strain (Fig. S3). From these data we conclude that Emp1 is required for macrophage cytolysis.

## Discussion

In this study, we demonstrated that Emp1, a predicted NAT, is required for the N-terminal acetylation of EsxA and other mycobacterial proteins. The orthologous gene from *M. tuberculosis, ERD_3144 (Rv2867)*, was also sufficient to restore the N-terminal acetylation of EsxA in the Δ*emp1* strain, supporting functional conservation between the two species. *In vivo*, Emp1 is solely responsible for the acetylation of EsxA and other mycobacterial proteins. In the Δ*emp1* strain, no acetylation of EsxA was observed (Fig. 3B). We demonstrated that Emp1 is dispensable for ESX-1-dependent secretion and hemolysis, and for growth in macrophages during infection. However, Emp1 was required for optimal macrophage cytolysis by *M. marinum*. Collectively, this study identified a NAT required for N-terminal acetylation in *Mycobacterium*, and provided insight into the requirement of N-terminal acetylation of EsxA and other proteins for mycobacterial virulence in the macrophage.

We previously identified and quantified N-terminal peptides in both *M. marinum* and *M. tuberculosis* (43). While ∼10-15% of the mycobacterial proteome is likely N-terminally acetylated (43), little is known about the NAT enzymes responsible for N-terminal acetylation in *Mycobacterium*. Prior studies aimed at understanding N-terminal acetylation have focused on EsxA. The initial study demonstrating N-terminal acetylation of EsxA suggested that EsxA acetylation impacted the interaction with its binding partner, EsxB (23). If this were the case, we would have expected a loss of EsxA and EsxB protein in the *Δemp1* strain, similar to the Δ*esxA* strain. Instead, EsxA and EsxB were made and secreted from *M. marinum* in the Δ*emp1* strain. Aguilera et al. mutated the Thr residue at the second position of EsxA, reporting reduced cytoplasmic translocation and macrophage cytolysis (25). They proposed that N-terminal acetylation of EsxA was required for ESX-1 function, suggesting that unacetylated EsxA was unable to disassociate from EsxB, preventing phagosomal lysis and macrophage cytolysis (25). In agreement with this study, abrogation of EsxA acetylation through the deletion of *emp1* did result in a significant reduction of macrophage cytolysis. However, we do not attribute the reduced cytolysis to a loss of ESX-1 function for several reasons. First, our prior work demonstrates that EsxA is required for ESX-1-dependent secretion and hemolytic activity because it is required for secretion of ESX-1 substrates (14, 66). Indeed, we have reported *M. marinum* strains that secrete EsxA and EsxB but are attenuated and non-hemolytic (14, 66, 73). If N-terminal acetylation was required for EsxA function, we would have expected a loss of protein secretion and hemolytic activity of the *Δemp1* strain. Instead, neither secretion nor hemolysis were dependent on Emp1. Second, in the absence of ESX-1 secretion, *M. marinum* is retained in the phagosome and is significantly attenuated of growth in the macrophage, similar to the Δ*eccCb*_*1*_ strain. If unacetylated EsxA resulted in a loss of EsxA function, we would have expected attenuated growth of the Δ*emp1* strain. Instead, growth of the Δ*emp1* strain during macrophage infection was comparable to the WT strain. We suspect changing the second residue of EsxA to modulate acetylation impacted the function of EsxA, resulting in a loss of secretion which would explain the lack of phagosomal damage, and the reduced cytolysis. In our study, the unacetylated EsxA protein retains its WT sequence and clearly promotes secretion and virulence. We suspect that the Emp1-dependent N-terminal acetylation of another protein or proteins is required for macrophage cytolysis, downstream of ESX-1 function. Alternatively, it could be that EsxA N-terminal acetylation contributes to cytolysis downstream of phagosomal lysis. Further work is required to distinguish between these two possibilities.

In this study, we advance the field of mycobacterial physiology by identifying a NAT that promotes N-terminal acetylation, contributing to the basic understanding of this fundamental protein modification in bacteria. We define a putative NAT required for EsxA acetylation, suggesting that N-terminal acetylation is dispensable for ESX-1 function under the conditions tested in this study, moving the field of Type VII secretion forward. We provide a framework for the identification of NATs required for the N-terminal acetylation of specific protein targets that is widely accessible and applicable to any system. Importantly, we showed that an antibody against an acetylated N-terminal peptide could discriminate between acetylated and unacetylated N-termini. The generation of similar antibodies for additional N-terminally acetylated proteins could be used in any system to demonstrate N-terminal acetylation and identify the responsible NAT.

Our study raises several questions about both N-terminal acetylation of mycobacterial virulence factors and the role of NATs in mycobacterial physiology and pathogenesis. First, it is unclear why EsxA is N-terminally acetylated or which Emp1 targets promote macrophage cytolysis. Second, Emp1 is required for a subset of the N-terminally acetylated mycobacterial proteins. It is unclear what dictates the specificity of Emp1, or what shared characteristics of proteins promote N-terminal acetylation by Emp1. Third, it remains unknown which of the additional 22 putative NATs contribute to N-terminal acetylation in *Mycobacterium* as well as their breadth of function and specificity.

One limitation of this study is that were we unable to show the Emp1 was sufficient and necessary for the acetylation of EsxA *in vitro*. This would support the hypothesis that Emp1 directly acetylates EsxA at its N-terminus. We expressed the Emp1 and the Emp1W223A versions in *E. coli* with the goal of purification from a heterologous host. Despite trying different temperatures, additives and vectors, we were unable to generate and isolate soluble forms of the proteins. Instead, we expressed Emp1 in *E. coli* and incubated the resulting lysate to acetylate a series of EsxA N-terminal peptides. Finally, we tried co-expressing *emp1* and either *esxA* or *esxBA* in *E. coli* and measuring EsxA acetylation using western blot analysis. We were unable to observe acetylation using these approaches. We are uncertain why we are unable to produce functional Emp1 protein *in vitro* or in *E. coli*, while we can express and purify functional NATs from *E*.*coli* and *S. typhimurium* (RimI). We suspect that Emp1 requires additional, unidentified cofactors or environmental cues for function that are specific to *Mycobacterium*.

Overall, this study contributes a fundamental understanding of the conserved mechanisms and underlying N-terminal protein acetylation in pathogenic mycobacteria and identifies the NAT solely responsible for EsxA acetylation in *M. marinum*, opening new avenues of study aimed at further understanding this protein modification in bacteria.

## Materials and Methods

*M. marinum* strains were grown as described previously (14). Plasmids were constructed using FAST Cloning or restriction cloning and maintained in *E. coli* as described (14, 73, 74). *M. marinum* strains were constructed using allelic exchange (14, 73, 74). Nt-Acetylation was measured using dot blot and NUT-PAGE followed by western blot analysis. Protein production and secretion were measured using western blot analysis as previously described (14). Site directed mutagenesis of the *emp1* gene was performed as in (56, 57). Whole colony MALDI mass spectrometry to measure surface associated EsxA, Nt-EsxA and EsxB was performed as in(58). Label free Quantitative Mass Spectrometry was used to measure Nt-acetylation from *M. marinum* whole cell lysates, similar to (68, 75). Hemolytic activity of *M. marinum* was measured against sheep Red Blood Cells (sRBCs) as previously described (14). Thin Layer Chromatography was used to confirm PDIM production in the Δ*emp1* strain (76). RAW264.7 cells were used as an infection model to measure growth of *M. marinum* during infection and macrophage cytolysis as described previously (14). Bioinformatic analysis and statistical analysis was performed using Prism and R Studio. Detailed materials and methods are available in the Supplementary Material.

## Acknowledgments

We would like to acknowledge the Mass Spectrometry and Proteomics and Genomics and Bioinformatics Facilities at The University of Notre Dame. SDW is funded from the Chemistry Biochemistry-Biology Interface (CBBI) NIH grant T32GM075762. We would like to thank Emily McMackin and Rachel Bosserman for the construction of the Δ*ppsC* and Δ*mas* strains used as controls in Figure S3. We thank the Champion Lab for the critical reading of this manuscript. P.A.C. is supported by the National Institutes of Health under award numbers AI156229, AI106872, AI149147, and AI149235.

