## Supplemental Figures and Tables for "An N-acetyltransferase required for EsxA N-terminal protein acetylation and virulence in *Mycobacterium marinum*"

**This PDF file includes:**

Supporting text (Materials and Methods)
Figures S1 to S3
Tables S1 to S2
SI References

**Other supporting materials for this manuscript include the following:**

Datasets S1

**Supplementary Materials and Methods**
**Maintenance of Bacterial Strains:**

All *M. marinum* strains were derived from the *M. marinum* M strain (ATCC BAA-535). *M. marinum* strains were maintained in Middlebrook 7H9 defined broth (Sigma-Aldrich, St Louis MO) supplemented with 0.5% glycerol and 0.1% Tween-80 (Fisher Scientific, Pittsburgh PA) or on Middlebrook 7H11 agar (Sigma-Aldrich) plates supplemented with 0.5% glycerol and 0.5% glucose. *M. marinum* was maintained at 30°C. When necessary, agar plates and broth were supplemented with 20 µg/mL kanamycin (IBI Scientific, Peosta, IA), 50 µg/mL hygromycin (EMD Millipore, Billerica, MA), or 60µg/mL X-gal (Millipore) as necessary. *E. coli* DH5α strains were grown at 37°C in LB (Luria-Bertani) media (VWR). When necessary, the following antibiotics were added to media at the following concentration: 50 µg/mL kanamycin, 200 µg/mL hygromycin, or 200 µg/mL ampicillin (Thermo Fisher, Waltham, MA).

**Protein Preparation and Analysis:**

<u>Secretion Assays.</u> ESX-1 protein secretion assays were performed as described previously (1). Briefly, *M. marinum* strains were grown in 7H9 media, diluted to an $OD_{600}$ of 0.8 in Sauton's Broth and grown for 48 hours. *M. marinum* was collected using centrifugation. The resulting supernatant was filtered through 0.2-µm Nalgene Stericups with polyethersulfone (PES) filters and concentrated by ultrafiltration in a 3,000-molecular-weight-cutoff Amicon filter (Millipore). Cell associated proteins were extracted from cells by lysis in PBS with a Biospec Mini-BeadBeater-24. Protein concentrations for cell-associated and secreted protein fractions were determined using the Pierce MicroBCA kit (Thermo Scientific).

<u>Western Blotting.</u> Unless otherwise noted, 10µg of protein were loaded onto 4-20% TGX-Gradient Gels (BioRad) for western blot analysis. All antibodies were diluted in 5% nonfat dry milk in PBS–0.1% Tween 20 at the following concentrations: RNAP polymerase

subunit β (RNAP-β) (ab12087; Abcam), 1:20,000; monoclonal ESAT-6 (EsxA, Thermo Fisher) 1:500 or polyclonal ESAT-6 (EsxA polyclonal antibody was produced by Genscript using the following peptide epitope: TEQQNFAGIEAAC), 1:1,000; Acetyl ESAT-6 (AcEsxA polyclonal antibody was produced by Genscript using the following peptide epitope: Acetyl-TEQQNFAGIEAAC) 1:1,000; polyclonal anti-*Mycobacterium tuberculosis* CFP10 (gene Rv3874) (antiserum, rabbit; 1:5,000) (NR-13801); polyclonal anti-*Mycobacterium tuberculosis* Mpt32 (gene Rv1860) (antiserum, rabbit; 1:20,000) (NR-13807). Horseradish peroxidase (HRP)-conjugated goat anti-mouse immunoglobulin secondary antibody (Bio-Rad) was utilized at 1:5,000 for detection of anti-RNAPβ and anti-EsxA (Thermo Fisher). HRP-conjugated goat anti-rabbit IgG secondary antibody (Bio-Rad) was utilized at 1:5,000 for detection with all antibodies.

NUT-PAGE: Neutral Urea Triton Polyacrylamide Gel Electrophoresis were performed as described in (2). NUT-PAGE resolving gels were composed of 6M Urea, 0.5M MOPs pH7, 4% Triton, 15% Polyacrylamide, 0.027% (w/v) APS, 0.13% (v/v) TEMED. Stacking gels were composed of 6M Urea, 9% Acrylamide, 100mM MOPs, 0.37% Triton-X, 0.06% (w/v) APS, 0.3% (v/v) TEMED. Running buffer was composed of 100mM Imidazole and 40mM MOPs pH7.10μg of protein were run in each lane for analysis. Gels were run for 1200V*hr, or 100V over the course of 12 hours. EsxA and AcEsxA were detected by Western Blotting as described above.

Dot Blot: 20μg of peptide were suspended in sterile water and applied to an Odyssey® nitrocellulose membrane and allowed to dry. Once completely dry (approximately one hour) were taken through western blotting with the following fluorescence-based reagents. The membrane was blocked using Blocking Buffer for Fluorescent Western Blotting (Rockland). Additionally, antibodies were diluted in this blocking buffer. The following primary antibodies were used simultaneously at the following dilutions: monoclonal ESAT-6 (EsxA, Thermo Fisher) 1:500 and 1:1,000; Acetyl

ESAT-6 (AcEsxA, GenScript). The following secondary antibodies were used simultaneously at the following dilutions: A 1:5,000 dilution of Goat Anti-Mouse IgG (H+L) DyLight$^{TM}$ 800 Conjugated (Thermo Scientific) and 1:20,000 dilution of Goat Anti-Rabbit IgG (H+L) DyLight$^{TM}$ 680 Conjugated (Thermo Scientific). Goat Anti-Rabbit was used to reveal Acetyl ESAT-6 (AcEsxA) and Goat Anti-Mouse was used to identify ESAT-6(EsxA). monoclonal ESAT-6(EsxA) (Thermo Fisher) 1:500 and 1:1,000; Acetyl ESAT-6 (AcEsxA) (GenScript). The following secondary antibodies were used simultaneously at the following dilutions: Goat Anti-Mouse IgG (H+L) DyLight$^{TM}$ 800 Conjugated (Thermo Scientific) and Goat Anti-Rabbit IgG (H+L) DyLight$^{TM}$ 680 Conjugated (Thermo Scientific). Goat Anti-Rabbit was used to reveal Acetyl ESAT-6(AcEsxA) and Goat Anti-Mouse was used to identify ESAT-6(EsxA).

**Generation of *M. marinum* Strains:**

Genetic deletions were constructed using allelic exchange as previously described (3-5). Upstream and downstream regions of genes of interest were amplified (roughly 1500 base pairs in both directions). These genetic regions were amplified with primers found in Supplementary Table 2, then introduced into the p2NIL vector (Addgene plasmid number 20188; a gift from Tanya Parish) by three-part FastCloning as previously described (6). Resulting constructs were digested with PacI (NEB) and dephosphorylated with Antarctic phosphatase (NEB). Enzymes were killed with heat treatment at 80°C. The pGOAL19 vector (Addgene plasmid number 20190; a gift from Tanya Parish) was digested with PacI followed by 65°C heat treatment. The resulting pGOAL was ligated into the p2NIL constructs. Plasmids were quantified on a NanoDrop instrument (Thermo Fisher), and 3 µg of plasmid was irradiated with 0.1 J/cm2 UV light in a CL-1000 UV cross-linker (UVP) followed by electroporation into 500µl of electrocompetent *M. marinum* cells using a GenePulser XCell (Bio-Rad). Electrocompetent cells were prepared exactly as described previously (7). Electroporated cells were recovered in 7H9 supplemented with

0.1% Tween 80 at 30°C overnight, spun down, and resuspended in 200µL of the same media for plating. Cells were plated on 7H11 (Sigma) supplemented with Kan, 60 µg/ml 5-bromo-4-chloro-3-indolyl-D-galactopyranoside (X-Gal), and oleic acid-albumin-dextrose-catalase (OADC). Merodiploids were picked, cultured, and then plated on 7H11 agar supplemented with OADC, 60 µg/ml X-Gal and 2% sucrose. White colonies were picked and cultured in 3 ml of 7H9 broth with 0.1% Tween 80. After approximately 5 days of growth, 500µl of culture was transferred to screw-cap tubes and 0.1 mm zirconia disruption beads (RPI) were added. Pellets were lysed by three 30 second pulses on a mini bead beater (Biospec Products) followed by a 10-minute centrifugation. 1µl of lysate was used in 10 µl PCR reactions. PCR products were run on TAE agarose gels stained with ethidium bromide (VWR) and imaged using a Gel Doc EZ imager (BioRad) and Image Lab software (BioRad). PCR products were interpreted as WT or knockouts based on the size of amplicons. Plasmids and primers used for generation of knock-out strains can be found in Supplementary Table 1 and 2.

**Generation of Complementation Plasmids:**

All complementation plasmids were generated by restriction digest using NdeI/SpeI (NEB) to isolate each gene from *puc19* plasmids containing genes of interest, followed by ligation of gene inserts into pMOP vector. All complementation plasmids were confirmed using targeted Sanger DNA sequencing at the Notre Dame Genomics and Bioinformatics Facility.

**Generation of Point Mutations Using Site Directed Mutagenesis (SDM):**

To generate the point mutation of Tryptophan at amino acid position 223 to an Alanine in the gene *MMAR_1839,* to generate a null mutant of the putative N-terminal acetyltransferase. SDM was performed as previously described, aligned with the instructions from the Quikchange II Site Directed Mutagenesis Kit (Agilent) (8-10). The primers are listed in Table S1.

**Hemolysis Assays:**

Hemolysis assays were performed as in Cronin et al (1), except that the hemolysis assay was incubated for 1.5 hours prior to reading. Briefly, *M. marinum* strains were grown in Middlebrook 7H9 broth (Millipore) and 0.1% Tween 80 (Thermo Fisher Scientific, Waltham, MA) to mid-log phase. The number of bacterial cells were normalized to absorbance at an optical density of 600 nm ($OD_{600}$). *M. marinum* cells were washed with phosphate buffered saline (PBS) and resuspended in 300 µl PBS. Sheep red blood cells (sRBCs, Hardy Diagnostics, Santa Maria, CA) were diluted 1 to 10 in PBS and subsequently washed PBS and gently centrifuged until supernatant was clear, removing dead sRBCs. sRBC were then resuspended in 500 µl PBS. Washed bacteria were mixed with 100 µl of sheep red blood cells, collected by centrifugation, and incubated at 30 °C for 1.5 hours. Each sample was read in technical triplicate in a 96-well plate on a SpectraMax ABS plate reader (Molecular Devices, San Jose, CA) at an optical density of 405 nm ($OD_{405}$). For all hemolysis assays, incubation of sheep red blood cells with water represents a maximal lysis positive control. Incubation of sheep red blood cells with PBS alone, or with the Δ*eccCb*$_1$ *M. marinum* strain, serve as negative controls for lysis. Error bars represent standard deviations. For each strain, data is representative of at least three biological replicates.

**Macrophage infections:**

<u>Cytolysis:</u> Cytolysis assays were performed similar to those in Bosserman et al (11). As described previously, RAW 264.7 murine macrophage cell line (ATCC TIB-71) was maintained in DMEM (Gibco) supplemented with 10% heat-inactivated fetal bovine serum (FBS, Seradigm) and maintained at 37°C with 5% $CO_2$. RAW 264.7 cells were plated at a density of $5 \times 10^5$ cells per well in cell culture treated 24-well plates (Greiner Bio-One) for 24 hours. Macrophages were infected with $2.5 \times 10^6$ *M. marinum* cell for a multiplicity of infection (MOI) of 5. Cells were incubated for 2 hours at 37 °C with 5% $CO_2$

and then treated with 100 μg/mL gentamycin (Research Products International, Mount Prospect, IL) for 4 hours at 37 °C to kill extracellular *M. marinum* cells. Monolayers were then washed three times with PBS and new media was added to each well. Macrophage staining was performed using Ethidium homodimer-1 and Calcein AM from the Live/Dead Viability/Cytotoxicity kit (Thermo Fisher Scientific, Waltham, MA) at 24 hours post-infection. Cells were imaged by Zeiss AxioObserver A1 inverted microscope using phase-contrast, rhodamine, and green fluorescent protein filters. At least three independent biological replicates, each in technical triplicate (3 wells), were performed for each strain. For each well, 10 image fields were captured. Dead cells were analyzed and counted using Image J as previously described (5).

CFUs: For CFU infections, as previously published in (11) except an MOI of 0.2 was used for *M. marinum* infections. The resulting *M. marinum* cells were plated at a dilution of 1:1000 for enumeration on agar. CFUs at the 2-hour timepoint were performed as described previously (11), except an MOI of 0.2 was used to keep MOI consistent with other timepoints, and cells were plated at 1:1000 dilution for enumeration on agar.

**Thin layer chromatography:**

Thin layer chromatography was performed as in (12) with the following changes. *M. marinum* strains were grown to saturation in Middlebrook 7H9 defined broth, and then diluted to an $OD_{600}$ of 0.05 in a 50ml volume. Following 48 hours of growth, the bacteria were collected and washed 3 times in PBS. Following lipid extraction, the samples were allowed to evaporate in a fume hood. Lipids were allowed to migrate via capillary action for approximately 15 minutes. PDIM lipids, specifically, were migrated 3 times sequentially. For PGL lipids, 6μl of lipid (the same volume of lipid that is used for PDIM) onto the TLC plates.

**Mass Spectrometry:**

Reagents and Equipment: LC-MS pure reagents (water, chloroform, acetonitrile, and methanol) were purchased from J.T. Baker (Radnor, PA). Tris(2-carboxyethyl)phosphine (TCEP) was purchased from TCI Chemicals (Portland, OR). Iodoacetamide (IAA) was purchased from VWR (Radnor, PA). S-Traps were purchased from Protifi. Unless specified, all other reagents were purchased from Sigma-Aldrich (St. Louis, MO). Hydrophilic–lipophilic balance (HLB) solid phase extraction (SPE) cartridges (10mg/1ml) were purchased from Waters Corporation (Milford, MA). Vacuum concentrator used was a Genevac miVac from SP Scientific (Warminster, PA).

LFQ: Sample preparation was performed similarly to as previously described (3, 11, 13). 100µg of protein from each *M. marinum* lysate samples were extracted by acetone precipitation. Dried samples were resuspended in 10µL 100mM TEAB, 5µL 1M TEAB, and 25µL 20% SDS. Proteins were reduced by addition of 5µL 100mM TCEP and incubated for 10 minutes at 95˚C. Proteins were subsequently alkylated with the addition of 5µL 100mM IAA for 30 minutes in darkness. Samples were acidified with 5µL of 12% $H_3PO_4$, and flocculated with 350µL of 90/10 methanol/100mM TEAB. Samples were each loaded and spun through S-Trap 'Minis' to immobilize proteins on-filter. Bound protein was washed with two additions of 150µL 90/10 methanol/100mM TEAB and one addition of 150µL 1:1 methanol/chloroform. Filters were placed into new collection tubes, and 2µg of sequencing grade trypsin (Promega, Madison, WI) in 160µL 100mM TEAB was added, to a 1:50 wt/wt enzyme-to-protein ratio. Samples were incubated for 4 hours at 37˚C. Following digestion, samples were spun down and eluted by two additions of 80µL 0.1% formic acid in water and one addition of 80µL 1:1 acetonitrile/water in 0.1% formic acid. Samples were dried on a vacuum concentrator for 20 minutes to reduce residual acetonitrile. Remaining sample was desalted with HLB SPE cartridges on a vacuum manifold according to manufacturer's instructions.

Samples were resuspended in 0.1% formic acid in water to 500ng/µL. Each sample had 1µL injected in triplicate into a Bruker nanoElute and timsTOF Pro MS system. A 90-minute 600nL/min gradient from 4-30% B (A= water + 0.1% formic acid, B= acetonitrile + 0.1 formic acid) was used on a 75µm x 100mm PepSep column with $C_{18}$ ReproSil AQ stationary phase at 1.9µm particle size, 120 Å pore size. CaptiveSpray nano-ESI with a spray voltage of 1700V. MS was set to PASEF mode with a mass range of 100-1700 m/z, ion mobility range of 0.6-1.6 v*s/cm$^2$, and ramp and accumulation times of 100ms. Each precursor consisted of 10 PASEF ramps for a cycle time of 1.17 seconds. Precursors were filtered to contain only charges from 2 to 5. MS/MS collision energy settings were set to ramp from 20eV at 0.6 ion mobility to 70eV at 1.6 ion mobility. Instrument tune parameters were set to default for proteomic studies with the following differences: quadrupole low mass set to 20 m/z, focus pre-TOF pre-pulse storage set to 5µs.

Peptide spectral mass matching and quantitative Label free quantification (LFQ) were performed on the raw data using PEAKS Online Xpro (build 1.4.2020-10-21_171258). A PEAKS Q analysis was performed to include label-free quantitative and FDR was controlled at the protein and peptide levels to <1%. (LFQ) data (14). The database searched against was an *M. marinum* M strain proteome FASTA file from Mycobrowser (Release 4). Fixed modification of carbamidomethylation of cysteines was set. Variable modifications were set: acetylation of protein N-terminus, deamidation of asparagine and glutamine, oxidation of methionine, conversion of glutamine and glutamic acid to pyroglutamic acid. Quantitation was normalized using the total ion current (TIC). All other search parameters were default unless specified. Raw and processed data are available through MassIVE and PrideDB ftp://MSV000091442@massive.ucsd.edu ftp://massive.ucsd.edu/MSV000091442/ Pride PXD040693.

Protein and peptide .csv files were exported from PEAKS, and the following analysis was performed using R. Each biological replicate had three strains: Wild type (WT), *ΔMMAR_1839,* and Complement (Comp), which each had three triplicate injections. Each peptide LFQ intensity was normalized to the protein RpoA LFQ value from its respective injection. For each biological replicate, the peptide was discarded if it was not observed in at least four observations from the six injections of WT and Comp (2 biological 3 technical replicates). Peptides were not considered if they did not start at the first or second canonical amino acid position of the protein. The average, standard deviation, and %RSD was calculated for each peptide for each strain. If there were not sufficient datapoints/degrees of freedom to calculate a standard deviation for a particular peptide/strain combination, the average %RSD for all other peptides in that peptide/strain combination was assigned. Peptides containing an N-terminal acetylation modification were separated from those that did not, ratios of average LFQ intensity for the *ΔMMAR_1839*/WT and Δ*MMAR_1839*/complemented were calculated from the non-terminal peptides. Peptides within proteins with ratios ≥ one were discarded. Remaining N-terminally acetylated peptides were matched with the non-acetylated cognate, and a total LFQ intensity for that peptide was calculated by summing the values. If multiple peptide variants existed for any protein (e.g. resulting from a missed cleavage or other PTM), the intensities of all peptides were summed together to obtain a total acetylated and non-acetylated LFQ value for each protein in the dataset. From these data % acetylation values were calculated by taking the ratio of acetylated LFQ intensity / total LFQ intensity for each protein. *k*-means clustering analysis was performed with each acetylated protein in the dataset. The variables considered for the clustering were the following LFQ intensity ratios: Δ*MMAR_1839*/Comp, Δ1839/WT, and Comp/WT, and clustering was performed with the kmeans() function in R, after each variable was centered using the base scale() function in R to reduce the influence of outliers. The proteins were clustered into three groups, using 25 random starting points, and those proteins clustering with EsxA were identified as potential acetylation targets of MMAR_1839. Code for this analysis can be found on GitHub (https://github.com/Champion-Lab/ESXA_Acetylation) along with a list of data analysis steps.

MALDI: Whole colony MALDI was performed identical to that described in Champion et al. (7). Briefly whole colonies from WT, Δ*eccCb*$_1$, Δ*MMAR_1839*, and the three complementation strains were picked and placed in a 0.45 µm spin filter (Costar) and briefly mixed with 100µl MS-grade water. Washate was collected by spinning and 1µl was spotted on a stainless steel MALDI target, dried and 1µl of a saturated solution of sinapinic acid in 50% acetonitrile/water was overlaid. MS spectra were acquired in linear mode on a Bruker Autoflex Extreme MALDI-TOF. A laser frequency of 500Hz was used and 10,000 spectra were summed with a mass range of 9,500 – 12,000 m/z. Mass calibration was performed using the $[M+1H]^{1+}$ and $[M+2H]^{2+}$ peaks from horse heart myoglobin spotted adjacent to the samples. MS spectra were exported as ASCII and imported into Prism Graphpad for visualization.

**Bioinformatics:**

The k-means clustering, generation of the SeqLogos, IceLogos and heatmaps were generated using R. All of the relevant code can be found on Github at the following link: https://github.com/Champion-Lab/ESXA_Acetylation.

**Statistical analysis:**

Statistical analysis was performed using Prism v. 9 as indicated in the figure legends.

```
EsxAMM          M-TEQQWNFAGIEAASSAIQGNVTSIHSLLDEGKQSLHKLAAAWGGSGSEAYRGVQQNWDS 60
EsxAMT          M-TEQQWNFAGIEAAASAIQGNVTSIHSLLDEGKQSLTKLAAAWGGSGSEAYQGVQQKWDA 60
                *-*************:********************* **************:****:**:

EsxAMM          TAQELNNSLQNLARTISEAGQAMSSTEGNVTGMFA 95
EsxAMT          TATELNNALQNLARTISEAGQAMASTEGNVTGMFA 95
                ** ****:***************:***********
```

**Fig. S1. Clustal alignment of the EsxA proteins from *M. marinum* and *M. tuberculosis.*** EsxAMM: MMAR_5450 from *M. marinum,* EsxAMT: Rv3875 from *M. tuberculosis.* Sequences were obtained from Mycobrowser. The N-terminal Met, green, is cleaved. The Thr at position 2, magenta, is the site of N-terminal acetylation. Shaded amino acids diverge between the two proteins.

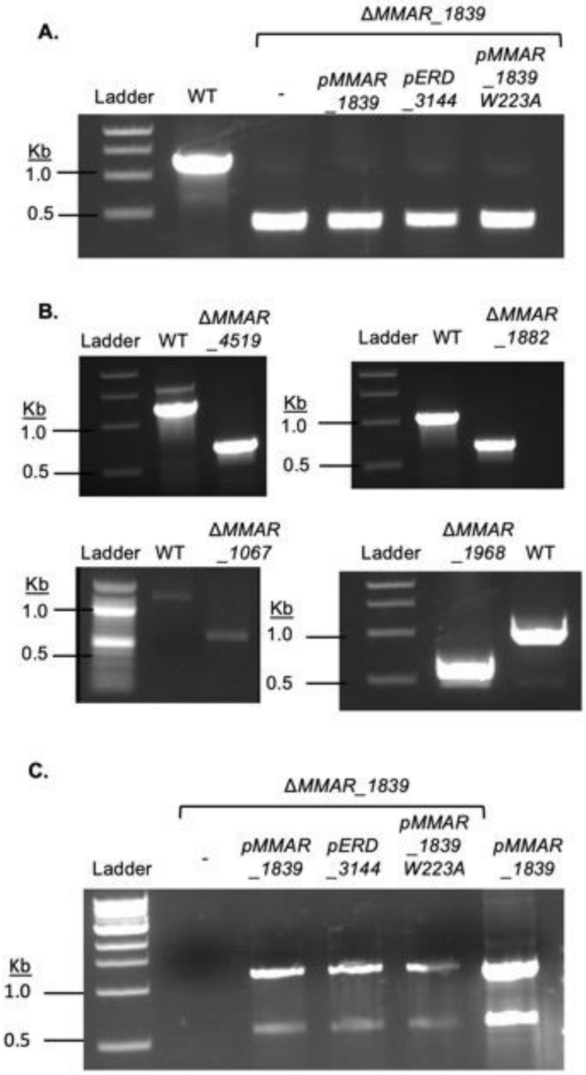


**Figure S2. Confirmation of Deletions and Complementation strains in *M. marinum*.** PCR confirming **A.** deletion of the *MMAR_1839* gene from *M. marinum* strains. Size of products in WT: 1218 bp and Δ*MMAR_1839*: 363bp. **B.** deletion of additional conserved predicted NAT genes. For *MMAR_4519,* WT: 1232 bp, Δ*MMAR_4519*: 578 bp, for *MMAR_1882,* WT: 1036 bp, Δ*MMAR_1882*: 565 bp, for *MMAR_1067*, WT: 1221 bp, Δ*MMAR_1067*: 555 bp, for *MMAR_1968,* WT: 1051 bp, Δ*MMAR_1968*: 526 bp. C. presence of the integrating expression plasmid for complementation in the Δ*MMAR_1839* strain. In the final lane, the pure plasmid was used as the template for the PCR reaction as a positive control. All PCR primers are listed in Table S2. Primers were designed to bind up and downstream from the predicted start based on annotation from Mycobrowser (15).

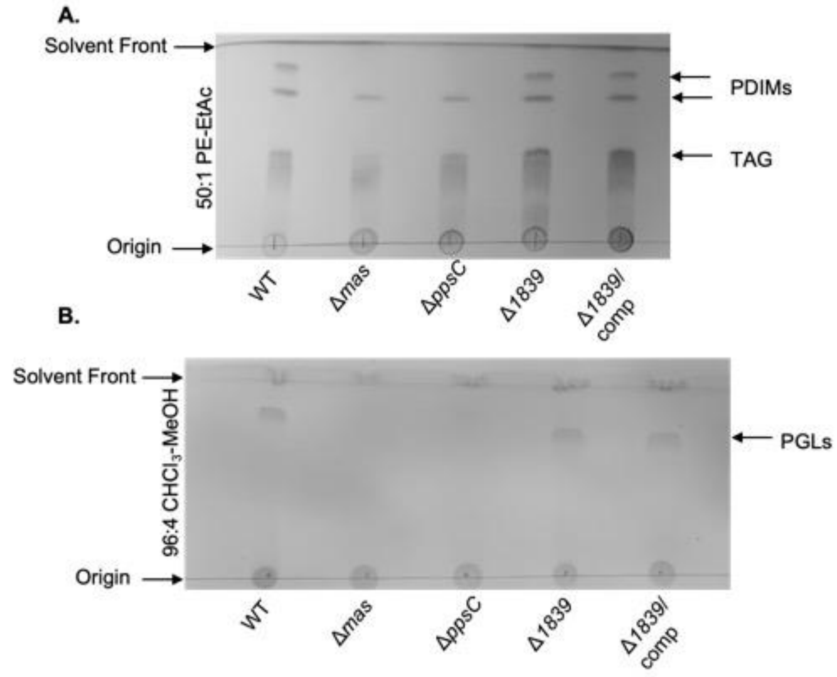


**Figure S3. TLC analysis of the relevant *M. marinum* strains.** Thin Layer Chromatography to confirm the presence of **A.** PDIMS and TAG, and **B**. PGLs. The WT strain serves as a positive control, the Δ*ppsC* and Δ*mas* strains serve as negative controls for all lipids measured. For all samples, 6μl of lipid extraction was spotted. The images are representative of at least three biological replicates.

**Table S1. Strains and plasmids used in this study**

| ***M. marinum*** **Strains** | | |
|---|---|---|
| **Name** | **Genotype** | **Reference** |
| *M. marinum* M strain | Wild-type strain; parental strain | ATCC BAA-535 |
| *ΔMMAR_1067* | M strain with an unmarked in-frame deletion of the *MMAR_1067* gene | This study |
| *ΔMMAR_1968* | M strain with an unmarked in-frame deletion of the *MMAR_1968* gene | This study |
| *ΔMMAR_4519* | M strain with an unmarked in-frame deletion of the *MMAR_4519* gene | This study |
| *ΔMMAR_1839* | M strain with an unmarked in-frame deletion of the *MMAR_1839* gene | This study |
| *ΔMMAR_1882* | M strain with an unmarked in-frame deletion of the *MMAR_1882* gene | This study |
| *ΔesxBA* | M with a deletion of the *esxBA* operon | (16) |
| *ΔMMAR_1839/ pMMAR_1839* | Δ*MMAR_1839* with the p*MMAR_1839* plasmid integrated at the *attB* site | This study |
| *ΔMMAR_1839/ pERD_3144* | Δ*MMAR_1839* with the p*ERD_3144* plasmid integrated at the *attB* site | This study |
| *ΔMMAR_1839/ pMMAR_1839W223A* | Δ*MMAR_1839* with the p*MMAR_1839W223A* plasmid integrated at the *attB* site | This study |
| *ΔppsC* | M strain with an unmarked in-frame deletion of the *ppsC* gene. | This study |
| *Δmas* | M strain with an unmarked in-frame deletion of the *mas* gene. | This study |
| **Plasmids** | | |
| p2NIL | parental suicide vector for allelic exchange, $Kan^R$, $Amp^R$ | (17), Addgene plasmid #20188 |
| pGOAL19 | marker cassette for allelic exchange, ampR, $Hyg^R$, *lacZ+ sacB* | (17), Addgene plasmid #20190 |
| p2NILΔ*1067*GOAL | Allelic exchange plasmid to generate the Δ*MMAR_1067* strain, contains *M. marinum* flanking regions (NC010612.1: 1289146...1290701;1291328…1292886) | This study |

| | | |
|---|---|---|
| | , p2NIL backbone with the GOAL19 marker cassette | |
| p2NILΔ*1968*GOAL | Allelic exchange plasmid to generate the Δ*MMAR_1968* strain, contains *M. marinum* flanking regions (NC010612.1: 2387011...2388617;2389075...2390645) , p2NIL backbone with the GOAL19 marker cassette | This study |
| p2NILΔ*4519*GOAL | Allelic exchange plasmid to generate the Δ*MMAR_4519* strain, contains *M. marinum* flanking regions (NC010612.1: 5547155...5548785;5549330...5550919) , p2NIL backbone with the GOAL19 marker cassette | This study |
| p2NILΔ*1839*GOAL | Allelic exchange plasmid to generate the Δ*MMAR_1839* strain, contains *M. marinum* flanking regions (NC010612.1: 2240745...2239051;2239899...2241609) , p2NIL backbone with the GOAL19 marker cassette | This study |
| p2NILΔ*1882*GOAL | Allelic exchange plasmid to generate the Δ*MMAR_1882* strain, contains *M. marinum* flanking regions (NC010612.1: 2292737...2294457;2294823...2296515) , p2NIL backbone with the GOAL19 marker cassette | This study |
| p2NILΔ*ppsC*GOAL | Allelic exchange plasmid to generate the Δ*ppsC* strain, contains *M. marinum* flanking regions (NC010612.1: 2156917...2158284;2159387...2160487) , p2NIL backbone with the GOAL19 marker cassette | This study |
| p2NILΔ*masGOAL* | Allelic exchange plasmid to generate the Δ*mas* strain, contains *M. marinum* flanking regions (NC010612.1: 2132139...22133557;2139863...2141313 ), p2NIL backbone with the GOAL19 marker cassette | This study |
| p*eccCb$_1$* | *eccCb$_1$* (MMAR_5456) expressed behind the mycobacterial optimal promoter(p$_{mop}$), pMH406 backbone; Hyg$^R$, parental plasmid for expression plasmids in this study | (12) |
| p*MMAR_1839* | p*MMAR_1839* expressed behind the mycobacterial optimal promoter(p$_{mop}$), integrated at *attB* | This study |

| | | |
|---|---|---|
| p*MMAR_1839W233A* | expressed behind the mycobacterial optimal promoter($p_{mop}$), integrated at *attB;* W residue at position 223 changed to an A | This study |
| p*ERD_3144* | p*ERD_3144* expressed behind the mycobacterial optimal promoter($p_{mop}$), integrated at *attB* | This study |

**Table S2. Oligonucleotides used in this study**

| Name | Sequence (5'-3') | Reference and Application |
|---|---|---|
| OMF241 | CGTGGTGTCACGCTCGTGGTGTCCATGCTGGCGTACC | PCR primers for amplifying flanking regions of *MMAR_1067* for deletion, underlined sequences match p2NIL, this study |
| OMF242 | GCAGCCTTAAGGTGGTCGTGGTCTGTCATGGC | |
| OMF243 | GACCACCTTAAGGCTGCCAACCCGGCCTGATCC | |
| OMF244 | ACGCAGTCAGGCACCGTCACGTTGTTCGAAGTCGATCAGCCAAAGG | |
| OMF251 | CGTGGTGTCACGCTCGTCGTTGTTGTACTCGACTGCCTTGG | PCR primers for amplifying flanking regions of *MMAR_1968* for deletion, underlined sequences match p2NIL, this study |
| OMF252 | GATGTTCTTAAGCCGGAAATCCTGTTGACTTTCG | |
| OMF253 | TCCGGCTTAAGAACATCTTGGGCAACACCCG | |
| OMF254 | ACGCAGTCAGGCACCGTTGGTACTCGACCGTGTCGTCG | |
| OMF259 | CGTGGTGTCACGCTCGTGGTGAAGTTCAGCTCACCGACGC | PCR primers for amplifying flanking regions of *MMAR_4519* for deletion, underlined sequences match p2NIL, this study |
| OMF260 | CACCGACTTAAGCAGCGGACCGACATCCATCG | |
| OMF261 | CGCTGCTTAAGTCGGTGGTCTCGACGCTGGTTCG | |
| OMF262 | ACGCAGTCAGGCACCGTCGCTCACTTGTGAACCGAGTCTGCG | |
| OMF267 | CGTGGTGTCACGCTCGTCGGTGACCTAACGGTCGAAGTGGGC | PCR primers for amplifying flanking regions of MMAR_1839 for deletion, underlined sequences match p2NIL, this study |
| OMF268 | GAAGGTCTTAAGCATGATGGGCGGAGCCGACATC | |
| OMF269 | TCATGCTTAAGACCTTCGCCACGGTGCTGCTCG | |
| OMF270 | ACGCAGTCAGGCACCGTCGTCCACCATCTTCGGGCTGATCG | |
| OMF275 | CGTGGTGTCACGCTCGTCAGCTTAGAGGCGATCACCGAAGC | PCR primers for amplifying flanking regions of MMAR_1882 for deletion, underlined sequences match p2NIL, this study |
| OMF276 | GGAATCTTAAGGGTTGGTACGTCAAGGTCTTTGGCC | |
| OMF277 | CCAACCCTTAAGATTCCGCACCTGCCCATGTTGC | |
| OMF278 | ACGCAGTCAGGCACCGTGTGGCCCGTCCATCGGTGAGC | |
| OGC3 | CCGGTCTGGAATCGTCAATC | |

| | | |
|---|---|---|
| OGC4 | CGCTGGGTCAGATTGGGATG | Δ*MMAR_1067* confirmation, this study |
| OGC5 | ATTCGGTGGCCATGCCGTAG | Δ*MMAR_1968* confirmation, this study |
| OGC6 | TGGTGGTGTCCAGTCAGTTC | |
| OGC7 | TGGGCAAACGCCAACCGATG | Δ*MMAR_4519* confirmation, this study |
| OGC8 | GCATGGGTACCAGCACAAAC | |
| OGC9 | CGTCAACGGTCCTGGTGAAG | Δ*MMAR_1839* confirmation, this study |
| OGC10 | AGAACAGTGACGCCAAGGAG | |
| OGC11 | TCAGCGAGTGCAGGTGATCC | Δ*MMAR_1839* confirmation, this study |
| OGC12 | TGTTCGCCATCGCGCAAGAC | |
| OGC25 | CAGATCCAGGGGGTTGCGGTTCATCCGGAGCGG | Site-directed mutagenesis primers for changing W to A at position 233 of MMAR_1839, changes from WT sequence underlined, this study |
| OGC26 | CCGCTCCGGATGAACCGCAACCCCCTGGATCTG | |
| OMF630 | ACTAGTCGGGACCGCTCAGGCGTCC | p*eccCb*$_1$ vector amplification for FAST cloning, (4). |
| OMF057 | CATATGGCTGGACTCCTGAATTCTGCAGCTG | |
| OMF096 | ACGAGCGTGACACCACGATGCC | p2NIL vector amplification for FAST cloning, (3) |
| OMF097 | ACGGTGCCTGACTGCGTTAGCAATTTAACTG | |
| oew264 | GTGGTGTCACGCTCGTTGGCCTCAAACGCGCATCAG | *ppsC*$_{ER}$ domain knockout, upstream |
| oew265 | CAAACCGCCCATGCCATTTACCACGGTGCGGCGAGATTCGAG | |
| oew266 | CGCACCGTGGTAAATGGCATGGGCGGTTTGGGTTTC | *ppsC*$_{ER}$ domain knockout, downstream |
| oew267 | GCAGTCAGGCACCGTTTGCCGTTTCCAGCTCGCTTG | |
| oew268 | AACGCTTTGTCCACCGACTG | Δ*ppsC*$_{ER}$ confirmation |
| oew269 | CGAGAAGGTCAGCCACCAATC | |

| | | |
|---|---|---|
| orb146 | TGGTGTCACGCTCGTGCAGTGAAGAACGGAGAGTC | *mas* knockout, downstream |
| orb147 | TCATAACGGCGGATCCGTTG | |
| orb148 | GGATCCGCCGTTATGAGGCAGGTTTTTGCATAAATC | *mas* knockout, upstream |
| orb149 | GCAGTCAGGCACCGTCGTTGTCGTTCTTGCCATTC | |
| orb159 | ACTGGATTCAGCCGGTGGTG | Δ*mas* confirmation |
| orb160 | ATCCGGCTTGGCCTGGATTG | |

**Dataset S1 (separate file).**

Tab S1A. Raw search results (.csv peptide/protein export from peptide-spectral mass matching.

Tab S1B. Trimmed Search Results Identical to SIA but contaminant/decoy proteins and unused columns have been removed.

Tab S1C. Bioreplicate Frequency. Trimmed, summed data for protein/peptide Termini for all N-terminally acetylated proteins observed across bioreplicates

Tab S1D. EsxA plot._Tabular data used to generate data in Figure 2

Tab S1E. Bioreplicate 3 cluster 3. Processed data as M&M and in Bioreplicate frequency used for k-means clustering in Figure 2. Shown are the predicted orthologs in *M. tuberculosis* H37Rv, protein product and functional categories derived from Mycobrowser(15).
